# Dopaminergic signals for reward, performance and social outcomes are dynamically gated during courtship

**DOI:** 10.1101/822817

**Authors:** Andrea Roeser, Vikram Gadagkar, Anindita Das, Pavel A. Puzerey, Brian Kardon, Jesse H. Goldberg

## Abstract

How does courtship affect dopaminergic (DA) responses to reward and motor performance outcomes? We used electrophysiology and fiber photometry to record DA signals in two mesostriatal pathways as thirsty male songbirds sang alone and to females. When alone, water reward signals were observed globally but singing-related performance error signals were restricted to a song-specialized mesostriatal pathway. During courtship singing, DA responses to both water-predicting cues and song performance outcomes diminished, and DA signals in the song pathway were instead driven by female calls timed with the male song. Thus DA signals are dynamically gated and routed through distinct pathways as animals change their priorities in response to a courtship opportunity.

---

Motivated behaviors are organized around singular objectives defined by an animal’s need^1^. For example, thirsty or hungry animals approach and learn from water- or food-predicting cues, in part due to reward prediction error (RPE) signaling by mesostriatal dopamine (DA) neurons^2, 3^. Recently, DA prediction error signals have also been observed in response to social and motor performance outcomes, supporting the generality of DA reinforcement mechanisms across a wide range of behaviors^4–8^. Yet it remains unclear how a single neuromodulatory system can evaluate outcomes in complex and natural conditions where an animal may weigh multiple objectives at once^9^. What if an animal is thirsty *and* sexually motivated *and* actively practicing motor performance? One possible solution is that a global DA signal is dynamically gated according to an animal’s current priorities. For example, DA responses to water cues would re- tune to social outcomes if an animal prioritizes courtship over thirst. A mutually inclusive possibility is that distinct mesostriatal DA pathways are anatomically segregated to signal outcomes according to distinct objectives^9^. This idea would predict that DA responses to social outcomes would not be globally broadcast but instead be specifically routed to striatal areas dedicated to social communication.

To test for dynamic gating, we examined how DA signals associated with three distinct objectives - quenching thirst, singing a good song, and courting a mate - change as thirsty, singing, and lonely male zebra finches were provided with opportunities to retrieve water, evaluate song performance errors, or court a female. Concurrently, to test if DA signals associated with distinct objectives are globally broadcast or anatomically routed to distinct striatal regions, we examined DA signals in two distinct mesostriatal pathways originating in the ventral tegmental area (VTA): one to Area X, the basal ganglia nucleus of a specialized song system^10^, and a different projection to a medial striatal area (MST) that arises from a distinct group of VTA DA neurons unassociated with communication circuits^11^.

We first examined behavioral and dopaminergic responses to water rewards. Water-deprived zebra finches were trained to peck, within eight seconds of a ‘reward’ light cue, a touch sensing spout that dispensed water with a probability of 70% (Methods). Birds also learned to ignore a different color ‘no-reward’ light cue unassociated with water (Retrieval rate following ‘reward’ and ‘no-reward’ light cues in lone, not-singing birds: 78.1±11.4% and 3.8±2.1%, respectively, *n* = 13 birds, Fig. 1a-c). We next used fiber photometry to measure DA release in Area X (*n* = 9 hemispheres, 7 birds) or MST (*n* = 9 hemispheres, 6 birds) (Fig. 1d, Methods). Both Area X and MST exhibited significantly larger activations of DA release following rewarded versus unrewarded spout contacts, consistent with classic RPE signaling (Extended Data Fig. 1)^2, 3^.

**Fig. 1.**
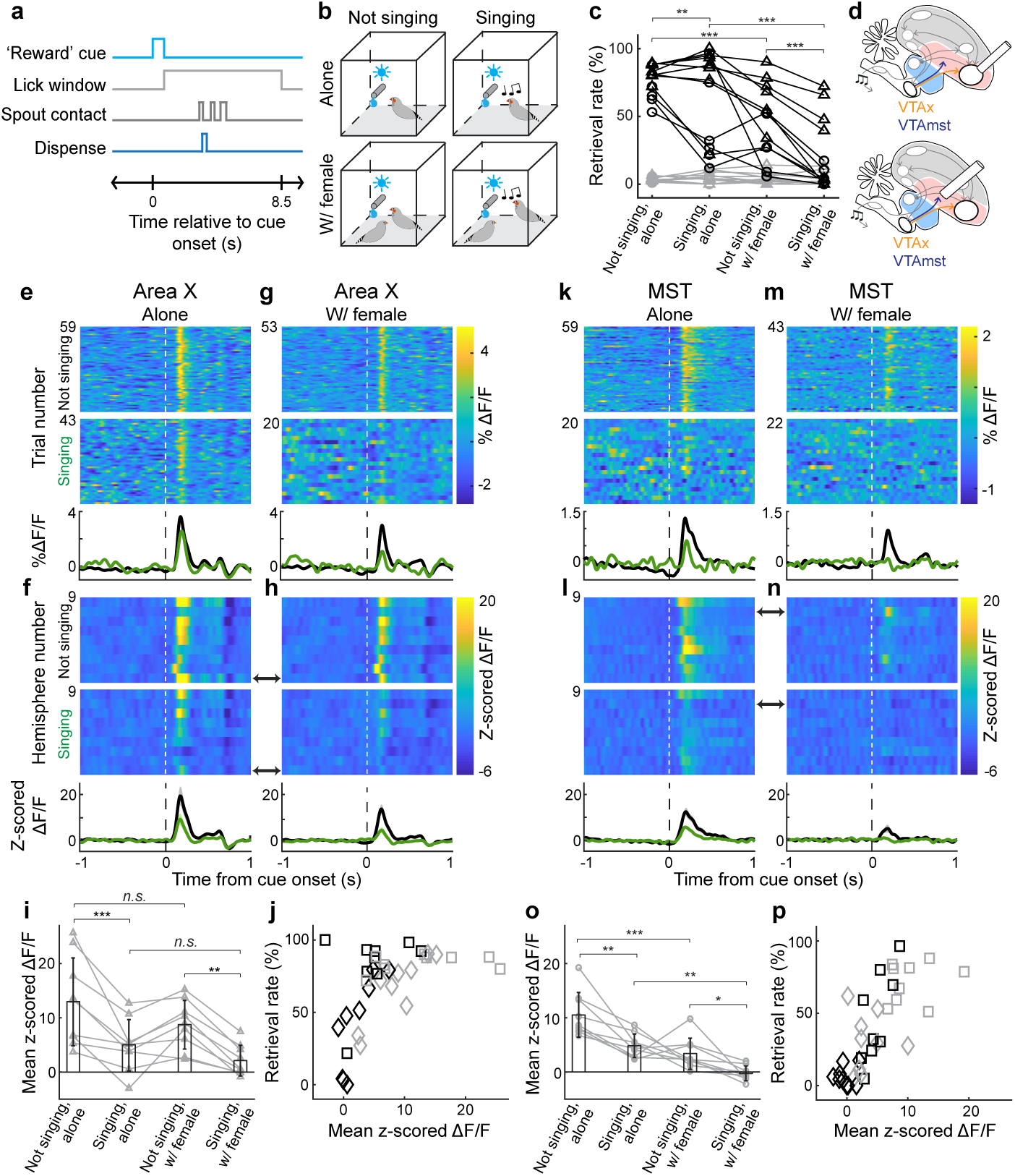
Courtship reduces behavioral and dopaminergic responses to water cues in water- deprived zebra finches. **a,** Example trial where ‘reward’ light cued water availability from a touch-sensing spout. **b,** Water availability under four behavioral conditions. **c,** Water retrieval probability for ‘reward’ (black) and ‘no-reward’ (gray) light cues across behavioral conditions (Area X implant: triangles; MST implant: circles). **d,** Brain diagrams indicating recording sites. **e- f,** DA responses when the bird was alone. **e,** Single trial DA responses to ‘reward’ light cue from a single Area X hemisphere during not singing (top) or singing (bottom) conditions. **f,** Average z- scored ΔF/F responses to ‘reward’ light cue across 9 Area X hemispheres (top), mean ± SEM z- scored ΔF/F across hemispheres (bottom). Black arrow denotes response from the example hemisphere in **e**. **g-h,** Data plotted as in **e-f** with female present. **i,** Average values across all hemispheres (mean ± SD, black) and mean z-scored values for each hemisphere (gray) for the four behavioral conditions in **e-h**. **j,** Retrieval rate in each behavioral condition plotted against the Area X DA response from that condition (black: singing; gray: not singing; square: alone; diamond: with female; R^2^ = 0.28 and *P*<0.001). **k-p,** Data plotted as in **e-j** for DA recordings in MST; **p,** R^2^ = 0.54 and *P*<0.001. * *P*<0.05, ** *P*<0.01, *** *P*<0.001, *n.s.* not significant for a 2-way ANOVA and post-hoc Tukey (**i** and **o**; Methods) and a generalized linear mixed effects model with post- hoc contrast tests (**c**; Methods); *n* = 9 Area X and *n* = 9 MST hemispheres.

To test if water retrieval depended on precisely what the animal was doing at the moment an opportunity to drink was provided, we presented light cues to water-deprived birds under four conditions: when they were alone and not singing, alone and singing, with the female and not singing, and with female and actively engaged in courtship song (Fig. 1b; Methods). Birds were most likely to retrieve water when alone and not singing, significantly less likely when singing, and least likely when singing to a female (Fig. 1c). We next measured DA responses to light cues in Area X and MST. Cue-evoked DA release was similarly robust and reliable in both Area X and MST when birds were alone and not singing (Fig. 1e,f,k,l, black), consistent with global responses to water-predicting cues in mammals^3^. Cue-evoked DA signals were significantly reduced when the male was singing alone, and reduced even more when singing to a female, consistent with the decreased water-seeking behavior during singing and courtship. Cue-evoked DA responses in both Area X and MST were significantly correlated with retrieval probability across conditions (Fig. 1j,p), showing that the reduction of DA signals by condition was global and associated with reduced interest in water. Yet interestingly, DA responses to ‘no-reward’ light cues also strongly diminished during both lone and courtship singing even though retrieval rates were always low, showing that the gating of DA signals can be decoupled from behavioral response (Extended Data Fig. 2). Altogether, these data show that the act of singing, especially to a female, reduces the expression of thirst and global DA responses to reward-predicting cues.

**Fig. 2.**
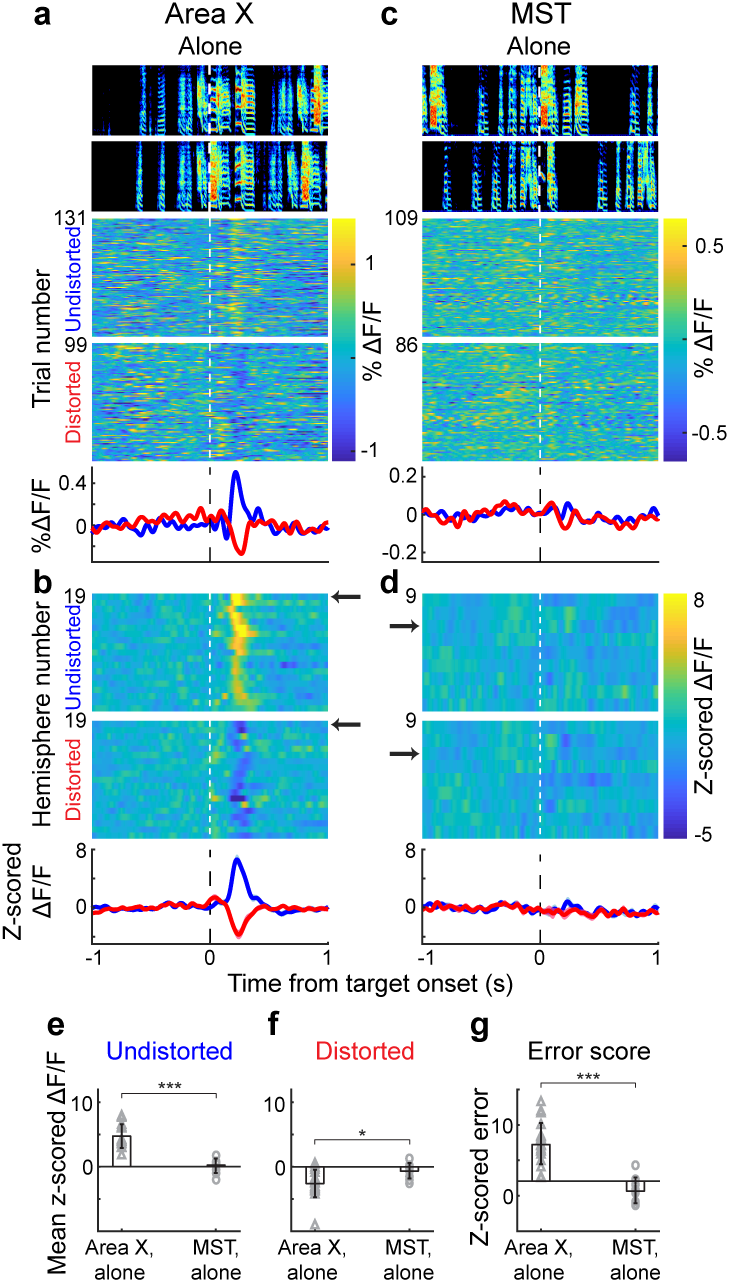
Singing-related error signals are routed to Area X and not MST in birds singing alone. **a,** Spectrograms and single trial DA responses for undistorted (top) and distorted (bottom) renditions recorded in a single Area X hemisphere of a bird singing alone, plotted above average % ΔF/F signals (all plots aligned to target onset; blue: undistorted; red: distorted). **b,** Z-scored average from 19 Area X hemispheres aligned to undistorted (top) and distorted (bottom) renditions, black arrow indicates example hemisphere shown in **a**; bottom: average z- scored response (mean ± SEM). **c-d,** Data plotted as in **a-b** for signals recorded in 9 MST hemispheres. **e-f,** Mean z-scored ΔF/F value for each hemisphere (gray) in the 150-300 ms window following undistorted (**e**) and distorted (**f**) renditions in Area X (triangles) versus MST (circles). Black: mean ± SD across all hemispheres. **g,** Z-scored error responses in Area X and MST for each hemisphere (gray) and across hemispheres (mean ± SD, black). Spectrograms in **a** and **c** (and all further instances) range from 0.5 - 7.5 kHz. * *P*<0.05, *** *P*<0.001 for an unpaired two-sided Wilcoxon rank sum test; *n* = 19 Area X and *n* = 9 MST hemispheres (**e-g**).

We next wondered, first, if singing-related DA signals were global or if they were routed to distinct striatal areas and, second, if they were gated by courtship state. When alone, zebra finches spontaneously engage in bouts of singing, their objective is to learn or maintain a song that matches the memory of a tutor song heard early in life^12^. Introduction of a female induces an immediate transition to a courtship state characterized by pursuit behavior and female-directed song^12, 13^. Past electrophysiological recordings from Area X projecting VTA neurons (VTAx neurons) in males singing alone discovered dopaminergic performance error signals: phasic activations in DA spiking followed better-than-predicted song outcomes, and phasic suppressions followed unexpected errors^4, 14^. To test if these singing-related error signals are globally broadcast or specifically routed to the song system, we measured DA release in Area X or MST in birds singing alone (Area X: *n* = 19 hemispheres, 14 birds; MST: *n* = 9 hemispheres, 6 birds). Perceived song errors were controlled with distorted auditory feedback (DAF), a 50 millisecond duration song-like sound played probabilistically on top of a specific target syllable in the bird’s song (Methods)^4, 10, 13, 15, 16^. In Area X, phasic suppressions of DA release followed DAF and phasic activations followed undistorted target syllable renditions at the precise moment that DAF would have occurred but did not (Fig. 2a,b), consistent with past electrophysiological recordings from VTAx neurons in lone males^4^. To quantify error responses, we compared the average DA fluorescence between distorted and undistorted songs in the 0.15-0.3 second interval following target onset (target onset defined as the median DAF onset time relative to distorted syllable onset, Methods). We defined the error score as the z-scored difference between target onset-aligned distorted and undistorted fluorescence^4, 17^, and found significant error signaling in all Area X hemispheres (error score >2 in 19/19 hemispheres; mean error score 7.4±2.9; Fig. 2g). In contrast, singing-related error signals were rarely observed in MST (error score >2 in 2/9 hemispheres; mean error score: 0.8±1.8; Fig. 2g). To our knowledge this is the first demonstration that DA performance error signals are not global and are instead routed to the specific part of the motor system producing the evaluated behavior.

We wondered if performance error signals in the VTAx pathway were dynamically gated by courtship state in the same way that water-reward signals were. If self-evaluation remains the goal during courtship singing, then error signals in the VTAx pathway might be similar with the female present or absent. Alternatively, if the bird’s objective changes, for example away from song evaluation and towards eliciting responses in the female, then DA signals may re-tune to respond to affiliative female behaviors. To test these possibilities, we measured DA responses to DAF-induced performance errors when birds sang alone and to females (Methods). Error signals that were robust in Area X when the male sang alone were reduced or gated off during courtship singing (percent reduction in phasic activations: 64.4±21.1%, *n* = 19 hemispheres; percent reduction phasic suppressions: 27.0±74.7%, *n* = 19 hemispheres; mean error score, alone = 7.4±2.9 vs. with female = 3.4±2.5, Fig. 3)^13^. To test if this gating of DA release in Area X also occurred in the spiking of VTAx DA neurons^18^, we carried out electrophysiological recordings from antidromically identified Area X projecting VTA neurons in singing birds hearing syllable-targeted DAF. We observed a similar courtship-associated reduction of performance error signals at the level of DA spikes (Extended Data Fig. 3, Methods). Again consistent with the anatomical routing of singing-related error signals, DA release in MST remained unresponsive to DAF during courtship singing (Extended Data Fig. 4).

**Fig. 3.**
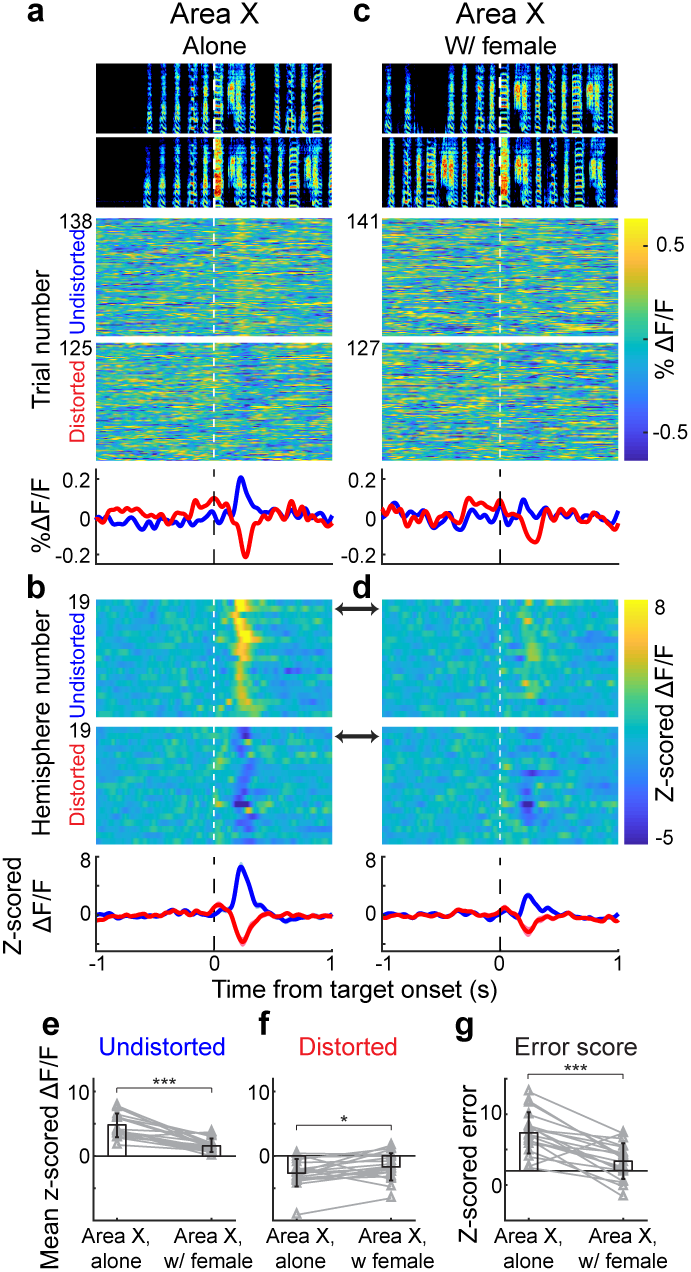
Singing-related error signals in Area X are reduced during courtship singing. **a,** Spectrograms and single trial DA responses for undistorted (top) and distorted (bottom) renditions recorded in a single Area X hemisphere of a bird singing alone, plotted above average ΔF/F signals (different example hemisphere from Fig. 2a; plots aligned to target onset; blue: undistorted; red: distorted). **b,** Z-scored average from 19 Area X hemispheres aligned to undistorted (top) and distorted (bottom) renditions, black arrow indicates example hemisphere shown in **a**; bottom: average z-scored response (mean ± SEM). **c,** Data plotted as in **a** for the same hemisphere measured during courtship singing. **d,** Data plotted as in **b** for the same hemispheres recorded during courtship singing. **e-f,** Scatter plots of average across all hemispheres (mean ± SD, black) and mean z-scored ΔF/F value for each hemisphere (gray) in the 150-300 ms following undistorted (**e**) and distorted (**f**) renditions in Area X during alone versus female directed singing. **g,** Z-scored error responses when birds sang alone versus to females (mean ± SD, black). * *P*<0.05, *** *P*<0.001 for a 2-way ANOVA and post-hoc Tukey (**e** and **f**; Methods) and for a paired two-sided Wilcoxon signed rank test (**g**); *n* = 19 Area X hemispheres.

**Fig. 4.**
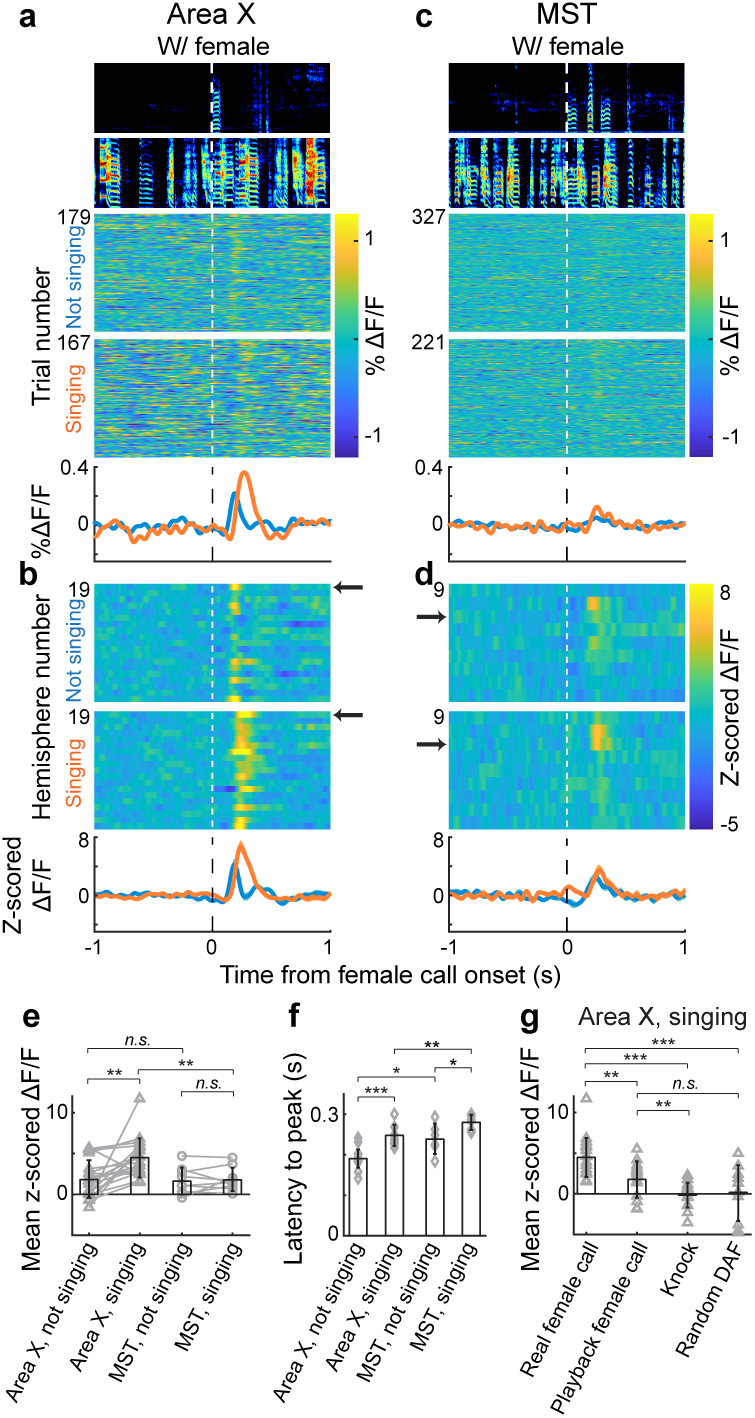
Female calls timed with male courtship singing drive phasic DA activations in Area X. **a,** Spectrograms and single trial Area X DA responses to natural female calls during not singing (top) or singing (bottom) periods of the courtship interaction, plotted above average % ΔF/F signals (plots aligned to onset of female call). **b,** Z-scored average from 19 Area X hemispheres, black arrow indicates example hemisphere shown in **a**; bottom: average z-scored response (mean ± SEM). **c-d** Data plotted as in **a-b** for DA signals recorded in MST. **e,** Scatter plots of average across all hemispheres (mean ± SD, black) and mean z-scored ΔF/F value for each hemisphere (gray) in the 150-300 ms window following female call onset produced during singing or not singing periods (triangles: Area X; circles: MST). **f,** Latency to peak z-scored response in a 300 ms window following real female calls during singing or not singing in Area X and MST (mean ± SD, black; only responses that were greater than a z-score value of 2 within the window were included). **g,** Z-scored magnitudes of DA responses to real female calls compared to playbacks of a female call, knock sounds, and random DAF sounds played through speakers next to the bird during courtship singing (mean ± SD, black; single hemispheres, gray). * *P*<0.05, ** *P*<0.01, *** *P*<0.001, *n.s.* not significant for a paired two-sided Wilcoxon signed rank test (within Area X and within MST in **e**) and for an unpaired two-sided Wilcoxon rank sum test (**f-g**; across Area X and MST in **e**); *n* = 19 Area X hemispheres and *n* = 9 hemispheres in MST.

We also noticed that baseline levels of DA in Area X increased during both alone and female-directed singing (Extended Data Fig. 5a-f), but this increase was not observed in the spiking of VTAx neurons (Extended Data Fig. 5m-r). And though the cued moment of female arrival drove phasic DA release, especially in MST (Methods; Extended Data Fig. 6e), baseline striatal DA levels did not increase with courtship state (Extended Data Fig. 6a-d and f). Baseline discharge of VTAx neurons during not-singing periods was subtly affected by courtship context (Extended Data Fig. 7i and j).

We hypothesized that gating of dopaminergic reward and performance evaluation signals during courtship allows the mesostriatal systems to re-tune to outcomes associated with a new objective, such as eliciting positive social feedback from the female. In zebra finches, calls provide affiliative feedback that supports bonding and reinforcement^12, 19^, suggesting that they may influence DA signals^20^. We aligned DA responses to natural female calls produced in two distinct phases of the courtship interaction: during periods when the male was not singing, and those produced during the male’s song (Fig. 4). Female calls produced when the male was not singing evoked small and unreliable DA release in MST (mean z-scored response: 1.7±1.7) and Area X (mean z-scored response: 1.9±2.3). In contrast, female calls produced during the male’s song drove dramatic and reliable DA release, but primarily in Area X (Area X mean z-scored response: 4.5±2.4; MST mean z-scored response: 1.9±1.4; Fig 4). To control for the possibility that any unexpected external sound during courtship song could drive DA signals, we played the sounds of knocks, DAF, or pre-recorded female calls throughout the courtship interaction (Methods). None of these stimuli drove reliable DA transients during singing (Fig. 4g, Extended Data Fig. 8). Thus only natural female calls in sync with the male song evoked DA release and specifically in Area X. To our knowledge this is the first demonstration that a temporally coordinated social interaction drives a phasic DA response specifically in a mesostriatal pathway dedicated to social communication.

A foundational premise of reinforcement learning (RL) is that animals learn to maximize future rewards^21^, but what an animal finds rewarding will depend on its current priorities. Indeed, recent studies showing DA responses to food and water rewards as well as social and motor performance outcomes support the generality of RL mechanisms^2–8^, but raise the question of how DA responses can evaluate diverse outcomes as an animal changes its objectives in natural, complex conditions. We discovered the DA system handles this problem in two ways. First, anatomically distinct mesostriatal DA pathways carried different signals: water reward signals were broadcast globally, but DA responses to song performance and social outcomes were routed specifically to a pathway dedicated to social communication. Second, DA signals re-tuned according to the current objective: both reward- and song performance-related DA signals were gated during courtship and were instead driven by female calls timed with male song. These results suggest that the bird’s evaluation system prioritizes social feedback during courtship above other objectives. It will be interesting to determine in future studies how these social feedback-driven DA signals influence future courtship interactions, pair formation, and mating success.

## Methods

### Animals

Subjects were 30 adult male zebra finches. All experiments were carried out in accordance with NIH guidelines and were approved by the Cornell Institutional Animal Care and Use Committee.

### Surgery - photometry

Birds were anesthetized with isoflurane and 1.25-1.75 uL of virus was injected either unilaterally or bilaterally into Area X (*n* = 12 right hemispheres; *n* = 7 left hemispheres; *n* = 14 birds) or MST (*n* = 3 right hemispheres; *n* = 6 left hemispheres; *n* = 6 birds) using a Nanoject II (Drummond) injector. Three different viral constructs were used: AAV9-CAG- GrabDA2m (*n* = 13 Area X hemispheres; *n* = 7 MST hemispheres; WZ Biosciences), AAV9-hSyn- GrabDA1h (*n* = 5 Area X hemispheres; *n* = 2 MST hemispheres; Addgene), AAV5-CAG- Dlight1.1 (*n* = 1 Area X hemisphere; Addgene). During the same surgery, optical fiber/s attached to a metal ferrule (Doric, 400 um core) were inserted above injection sites (Right/left hemisphere implants—Area X: +5.6A, +/-1.5L [relative to lambda], 2.5V [relative to pial surface]; MST: +4.7A, +/-0.7L, 2.9V; head angle 20 degrees). Data were collected at least 8 weeks post-surgery to allow for viral expression.

### Surgery - electrophysiology

During implant surgeries, birds were anesthetized with isoflurane and a bipolar stimulation electrode was implanted into Area X at established coordinates (+5.6A, +1.5L [relative to lambda], 2.65V [relative to pial surface]; head angle 20 degrees)^4^. All VTAx neurons in this dataset are from different birds than those reported in^4^. For recording VTAx neurons, custom microdrives carrying an accelerometer, linear actuator, and homemade electrode arrays (5 electrodes, 3-5 MOhms, microprobes.com) were implanted into a region where antidromically identified VTAx neurons were intraoperatively identified (*n* = 4 birds). For recording VTAother neurons, either custom microdrives as described above (*n* = 4 birds) or 16- channel movable electrode bundles (Innovative Neurophysiology) were used (*n* = 3 birds).

### Water reward training

Birds acclimated to the homecage for 2 to 3 days with ad lib food and water, a mirror, a perch, an inactive water spout, and distinct red and white LED lights. During the first phase of training, ad lib water was removed and the ‘reward’ light cue (0.5 second illumination of red or white LED light; ‘reward’ light color varied by bird) was presented with an exponentially distributed, average inter-trial interval of 180 seconds. At the offset of illumination, 5.0±1.5 uL of water was dispensed from the spout below the ‘reward’ light with a 100% water reward probability. After birds learned to reliably peck the spout following the ‘reward’ light cue (greater than 70% retrieval rate), we made water dispensation contingent on spout contact within 8 seconds of the ‘reward’ light cue offset. Next we introduced a distinct 0.5 second ‘no-reward’ light cue of different color on the opposite side of the homecage from the ‘reward’ light cue; it did not have a spout underneath, and never resulted in a water reward. ‘Reward’ and ‘no-reward’ light cue trials were randomly interleaved and presented at exponentially distributed inter-trial intervals averaging 150 seconds. Once birds learned to ignore the ‘no-reward’ cue (less than 10% retrieval rate), while still maintaining a greater than 70% ‘reward’ cue retrieval rate, head-mounted optical ferrules were attached to a fiber optic cable. Two to three days before optical imaging, the ‘reward’ light cue water reward probability was changed from 100% to 70%. To test if water retrieval rate is influenced by singing, female presence, and their interactions, we fitted generalized linear mixed models with binomial distribution using the *glmmTMB* package in *R*. We then conducted post-hoc contrast tests between conditions using the *emmeans* package in *R*. P-values were corrected for multiple testing using the Tukey method (Fig. 1c and Extended Data Fig. 2c).

### Syllable-targeted distorted auditory feedback

Postoperative birds were placed in a sound isolation chamber equipped with a microphone and two speakers which provided distorted auditory feedback (DAF). To implement targeted DAF, the microphone signal was analyzed every 2.5 ms using custom LabVIEW software. Specific syllables were targeted by detecting a unique inter-onset interval (onset time of previous syllable to onset time of target syllable) using the sound amplitude as previously described^4^. The targeted syllable was programmed to be distorted with DAF 50% of the time. DAF was a broadband sound bandpassed at 1.5-8 kHz– the same spectral range of zebra finch song. DAF amplitude was measured with a decibel meter (CEM DT-2 85A) and maintained at less than 90 dB.

### Photometry data acquisition and analysis

470 nm and 405 nm LEDs (Doric LED Driver and Doric Connectorized LEDs) were modulated sinusoidally at 208.6 and 530.5 Hz respectively using custom LabVIEW code. Excitation signals were passed through minicube filters (Doric Fluorescence MiniCube) to the bird through a commutator (Doric Pigtailed Fiber-optic Rotary Joint). Emission signals were measured using a femtowatt photodetector (Newport, Model 2151) or an integrated 4 port minicube and photodetector (Doric) at 40kHz (to match microphone sampling rate). Demodulation with custom LabVIEW code produced a 470nm (DA) and a 405 (control) signal which were used to calculate the percent change of the fractional fluorescence signal (%ΔF/F=100*(470-405_fit)/mean[470]). Phasic signals were high-pass filtered (4 Hz butterworth); baseline signals at transitions to singing (Extended Data Fig. 5a-l) and courtship state (Extended Data Fig. 6a-d,f) were low-pass filtered (0.25 Hz butterworth). Hemispheres were excluded if peak fluorescence response across all experiments was less than a z-score of 2 or if fiber placement missed Area X or MST. Z-scored ΔF/Fs were calculated as (%ΔF/F-mean[baseline %ΔF/F]) / std[baseline %ΔF/F]. Baselines were defined as the one second preceding song target, light cue, or spout contact (Figs. 1-4 and Extended Data Figs 1, 2, 4, 6, and 8). Here and elsewhere in the paper, for analysis of transitions to singing from not-singing periods, singing onset was defined as the first of a sequence of at least five syllables with a maximum inter-syllable interval of 0.5 seconds; singing offset was defined as the offset of the last syllable of a song sequence followed by at least 0.5 seconds of silence. Accordingly, a 0.5 second baseline was used to ensure that the baseline not-singing window excluded song syllables (Extended Data Fig. 5). To quantify phasic responses to DAF, female calls, reward cues and knocks, we calculated the mean z-scored ΔF/F in a 0.15-0.3 second window after event onset (Figs. 1-4, Extended Data Figs. 2, 4, and 8). Due to a wider range of latencies to responses to spout contacts (Extended Data Fig. 1), a 0.17-0.5 second window was used to quantify RPE signals. Latencies for significant phasic signals were analyzed and quantified as the interval between event onset and peak (or dip) in the windows specified above (Fig. 4 and Extended Data Figs. 1 and 8). To test for change in baseline DA levels at transitions from not-singing to singing (Extended Data Fig. 5), we compared the mean z-scored ΔF/F in the 0-0.5 second window before and after song onsets (or offsets). To test for baseline changes in DA release before versus after female appearance (Extended Data Fig. 6), we compared the mean z-scored ΔF/F in the 1 second before her cued arrival to the 1-2 second interval after (to allow for the phasic signal to subside). To test the effects of syllable distortion or rewards, female presence, and their interactions on DA release, we used a within-subject two-way ANOVA model with the *aov* function in *R*. We then conducted post-hoc contrast tests between conditions using the *emmeans* package in *R*. P-values were corrected for multiple testing using the Tukey method (Fig. 3e-f, Extended Data Fig. 4 e-f, Fig 1i and o, Extended Data Fig. 2i and n). Unpaired two-sided Wilcoxon rank sum tests were used to test for significance across different groups of hemispheres (Fig. 2e-g, Fig. 4 e-g, Extended Data Fig. 1e-f, Extended Data Fig. 6e, and Extended Data Fig. 8 e,j,k,p, and q). Paired two-sided Wilcoxon signed rank tests were used to test for significance across the same hemispheres (Fig. 3g, Fig. 4e, Extended Data Fig. 1e, Extended Data Fig. 4g, Extended Data Fig. 5 e,f,k,l,q and r, and Extended Data Fig. 8e, j, and p). We used a one-sample t test to compare baseline DA levels before versus after female appearance (Extended Data Fig. 6f).

### Electrophysiology data acquisition and analysis

For the custom microdrives, neural signals were band-passed filtered (0.25-15 kHz) in homemade analog circuits and acquired at 40 kHz using custom Matlab software. For the 16-channel movable electrode bundles (Innovative Neurophysiology), neural recordings were obtained using 16-channel INTAN headstages with the accompanying recording controller and INTAN acquisition software at a sampling rate of 20 kHz. Single units were identified as Area X projecting (VTAx) by antidromic identification (stimulation intensities 50-400 μA, 200 μs on the bipolar stimulation electrode in Area X), as previously described^4^. Neurons not identified as projecting to Area X were defined as VTAother neurons. Spike sorting was performed offline using custom MATLAB software. Instantaneous firing rates (IFR) were defined at each time point as the inverse of the enclosed interspike interval (ISI). Firing rate histograms were constructed with 25 ms bins and smoothed with a 3-bin moving average. To calculate the mean rate and median ISI during singing (Extended Data Fig. 7), the firing rate and median ISI were averaged over all song motifs, with a time-window extending 50 ms before motif- onset to 50 ms after motif-offset. The coefficient of variation (CV) of the ISI and the peak of the spike-train autocorrelation (STA) in Extended Data Fig. 7 were computed over the entire singing bouts. To test for error responses, we compared the firing activity between randomly interleaved undistorted and distorted song renditions. We computed the z-scored difference between the target time-aligned distorted and undistorted firing rate histograms (Extended Data Fig. 3). The target time was defined as the median DAF onset-time relative to the distorted syllable onset-time. The error response was defined as the mean z-scored difference in a 50-125 ms window following target time^4^. To calculate the significance bars shown in Extended Data Fig. 3, spiking activity within ±1 second relative to target onset was binned in a moving window of 30 ms with a step size of 2 ms. Each bin after the target time was tested against all the bins in the previous 1 second (the prior) using a z-test^4^. To test the effects of syllable distortion, female presence, and their interactions on DA spiking, we used a within-subject two-way ANOVA model with the *aov* function in *R*. We then conducted post-hoc contrast tests between conditions using the *emmeans* package in *R*. P-values were corrected for multiple testing using the Tukey method (Extended Data Fig. 3 e-f). A paired two-sided Wilcoxon signed rank test was used to test for significance in Extended Data Fig. 3g, Extended Data Fig. 5 w and x, and Extended Data Fig. 7.

### Courtship interactions

Experiments were performed in the male’s homecage in a sound isolation chamber. During electrophysiological recording sessions, spiking data were first collected when the male was singing alone for at least 35 song motifs before the female was introduced. For fiber photometry experiments where we examined DA release at the transition to courtship state (Extended Data Fig. 6), we controlled for the precise moment of perceived female arrival by playing two female calls through speakers immediately prior to presenting the female. In all experiments, the female cage was placed next to the male’s within the sound isolation chamber allowing birds to hear and see each other.

### Data availability

The data that support the findings of this study are available from the corresponding author upon request.

## Acknowledgments

We thank Ali Mohebi, Nao Uchida, Brendan Ito, and the Goldberg lab members for comments, Arnav Raha for artwork, Zhilei Zhao for statistical advice, and Ariana Enzerink and Archana Podury for technical assistance. VG was supported by a Simons Foundation Postdoctoral Fellowship and a NIH/NINDS Pathway to Independence Award (grant # K99NS102520), PAP by NIH/NINDS (grant # F32NS098634), and JHG by NIH/NINDS (grant # R01NS094667).

## Author Contributions

AR, VG and JHG designed the research, analyzed data, and wrote the paper. AR, VG, AD, PAP and BK performed experiments.

---

**Extended Data Fig. 1.**
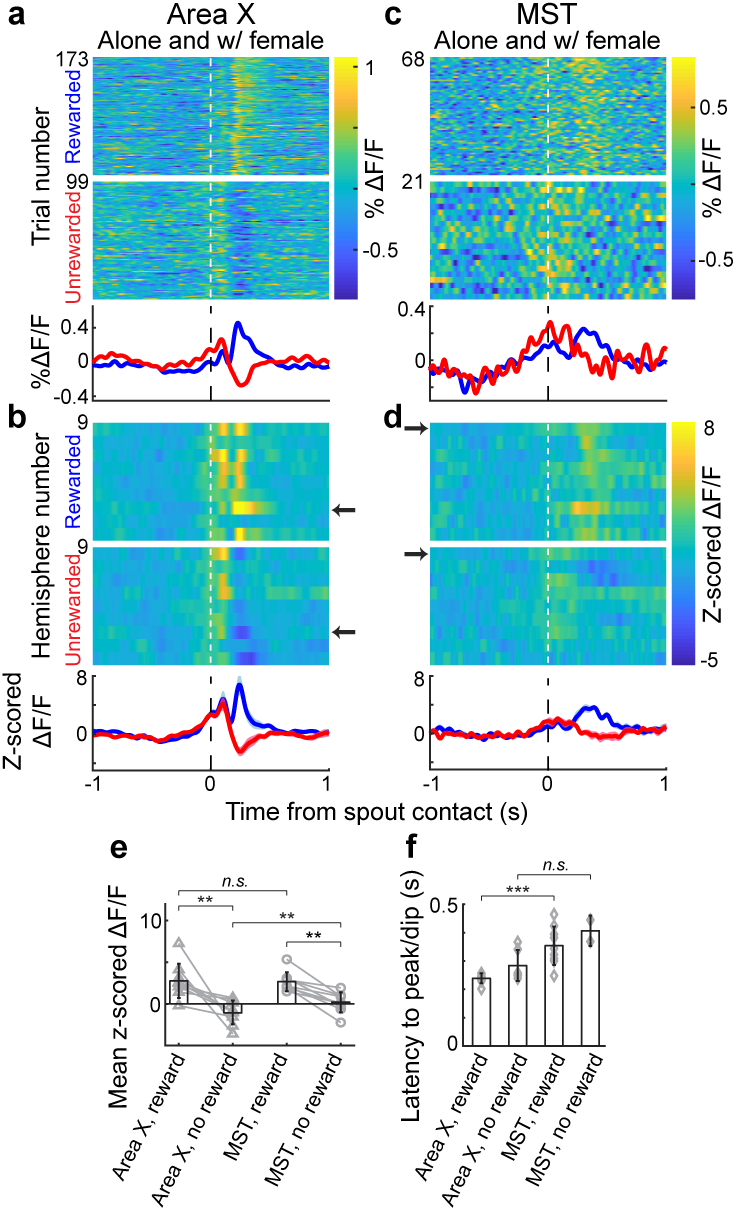
Reward prediction error signals exist in Area X and MST. **a,** Single trial DA responses from one Area X hemisphere in response to a water reward (top) or reward omission (bottom), aligned to spout contacts following ‘reward’ light cues. **b,** Average z-scored % ΔF/F signals across 9 Area X hemispheres, example hemisphere in **a** indicated by black arrow (top), average across all hemispheres (mean ± SEM; bottom). **c-d,** Data plotted as in **a-b** for MST. **e-f,** Reward signals were larger and faster in Area X compared to MST. **e,** Scatterplots showing average values across all hemispheres (mean ± SD, black) and mean z-scored values for each hemisphere (gray) for reward and reward omission in the 170-500 ms window following spout contact. **f,** Latency to peak (rewarded trials) or dip (omission trials) of the z-scored response in the 170-500 ms window after spout contact (mean ± SD, black; single hemispheres, gray; only responses that were greater than a z-score value of 2 within the window were included). ** *P*<0.01, *** *P*<0.001, *n.s.* not significant for a paired two-sided Wilcoxon signed rank test (within Area X and within MST in **e**) and for an unpaired two-sided Wilcoxon rank sum test (**f**; across Area X and MST in **e**); *n* = 9 Area X hemispheres and *n* = 9 hemispheres in MST.

**Extended Data Fig. 2.**
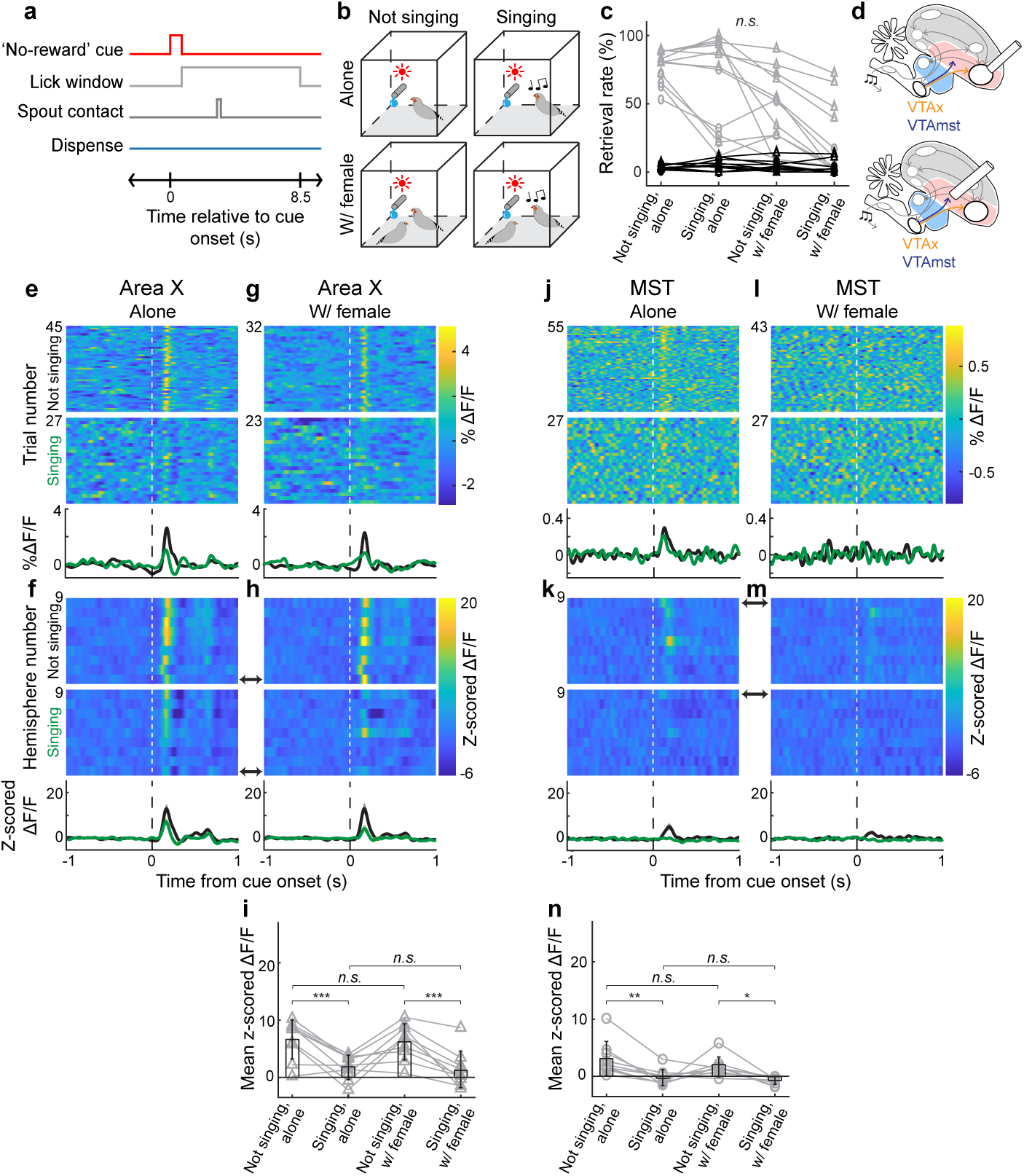
Singing reduces dopaminergic responses to ‘no-reward’ light cues in both Area X and MST. **a,** Example ‘no-reward’ light cue trial. **b,** Water availability under four behavioral conditions. **c,** Water retrieval probability for ‘no-reward’ (black) and ‘reward’ (gray) light cues across behavioral conditions (Area X implant: triangles; MST implant: circles). **d,** Brain schematics showing recording sites. **e-f,** DA responses to the ‘no-reward’ light when the bird was alone. **e,** DA responses for single trials from a single Area X hemisphere during not singing (top) or singing (bottom) conditions. **f,** Average z-scored ΔF/F signals across 9 Area X hemispheres, example hemisphere in **e** indicated by black arrow (top), average across all hemispheres (mean ± SEM; bottom). **g-h,** Data plotted as in **e-f** when males were with a female. **i,** Average values across all hemispheres (mean ± SD, black) and mean z-scored values for each hemisphere (gray) for the four behavioral conditions in **e-h**. **j-n,** Data plotted as in **e-i** for DA recordings in MST. * *P*<0.05, *** *P*<0.001, *n.s.* not significant for a 2-way ANOVA and post-hoc Tukey (**i** and **n**; Methods) and a generalized linear mixed effects model with post-hoc contrast tests (**c**; Methods); *n* = 9 Area X and *n* = 9 MST hemispheres.

**Extended Data Fig. 3.**
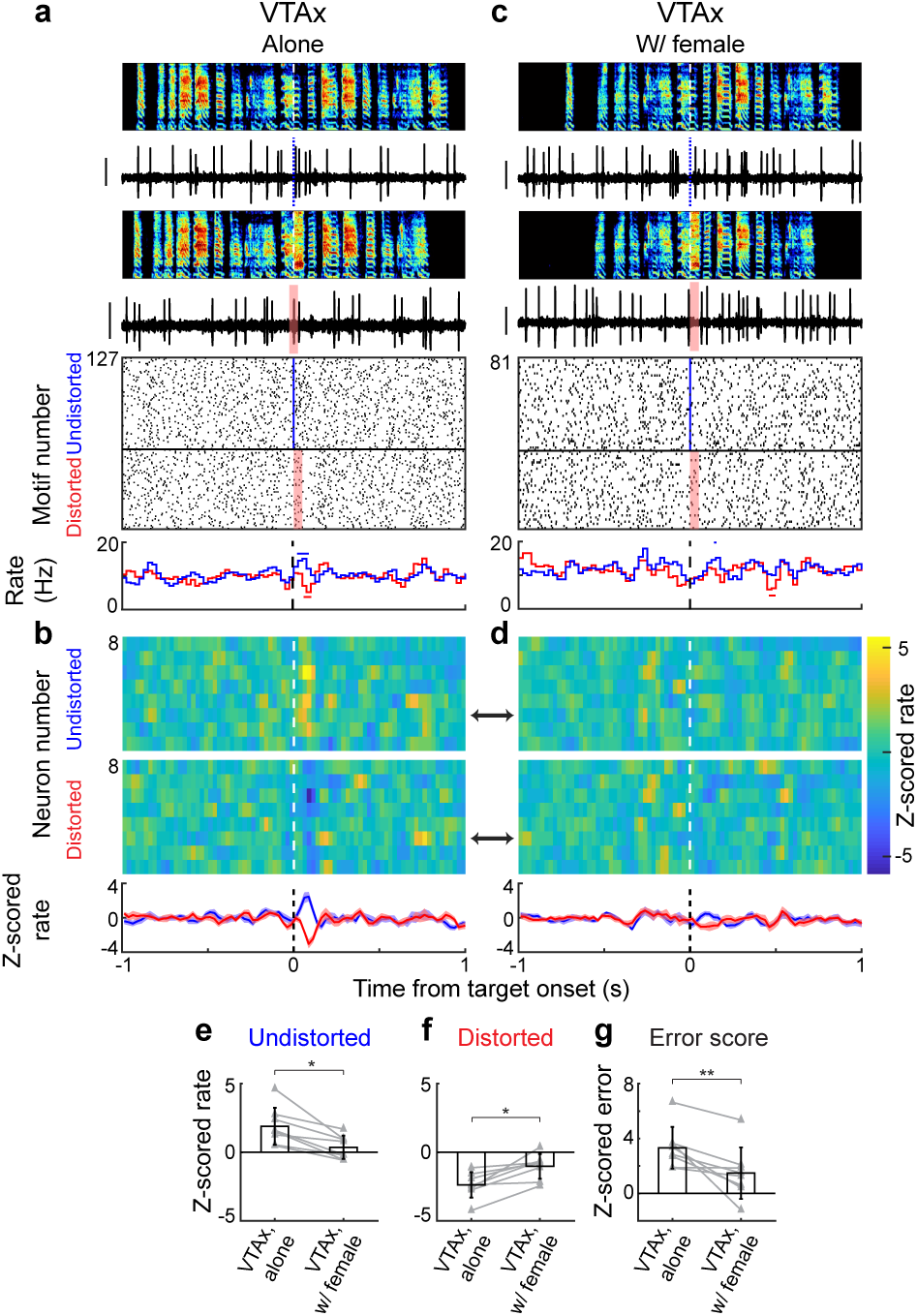
Gating of singing-related error signals during courtship is also observed at the level of VTAx spikes. **a,** Spectrograms, single trial spiking activity, and raster plots for undistorted (top) and distorted (bottom) renditions recorded in a single antidromically identified VTAx neuron of a bird singing alone, plotted above firing rate histograms (plots aligned to target onset; blue: undistorted; red: distorted). **b,** Z-scored average from 8 antidromically identified VTAx neurons aligned to undistorted (top) and distorted (bottom) renditions, black arrow indicates example neuron shown in **a**; bottom: average z-scored response (mean ± SEM). **c,** Data plotted as in **a** for the same VTAx neuron recorded during courtship singing. **d,** Data plotted as in **b** for the same VTAx neurons recorded during courtship singing. **e-f,** Scatter plots (gray) and mean ± SD (black) of average z-scored firing rate in the 50-125 ms following undistorted (**e**) and distorted (**f**) renditions during alone versus female directed singing across all VTAx neurons. **g,** Z-scored error responses for each neuron (gray) and mean ± SD (black) when birds sang alone versus to females. Horizontal bars in histograms (**a, c**) indicate significant deviations from baseline, *P*<0.05 for a one-sided z test. Scale bars in **a** and **c** for spiking activity is 1 mV. * *P*<0.05, ** *P*<0.01 for a 2-way ANOVA and post-hoc Tukey (**e** and **f**; Methods) and for a paired two-sided Wilcoxon signed rank test (**g**); *n* = 8 VTAx neurons.

**Extended Data Fig. 4.**
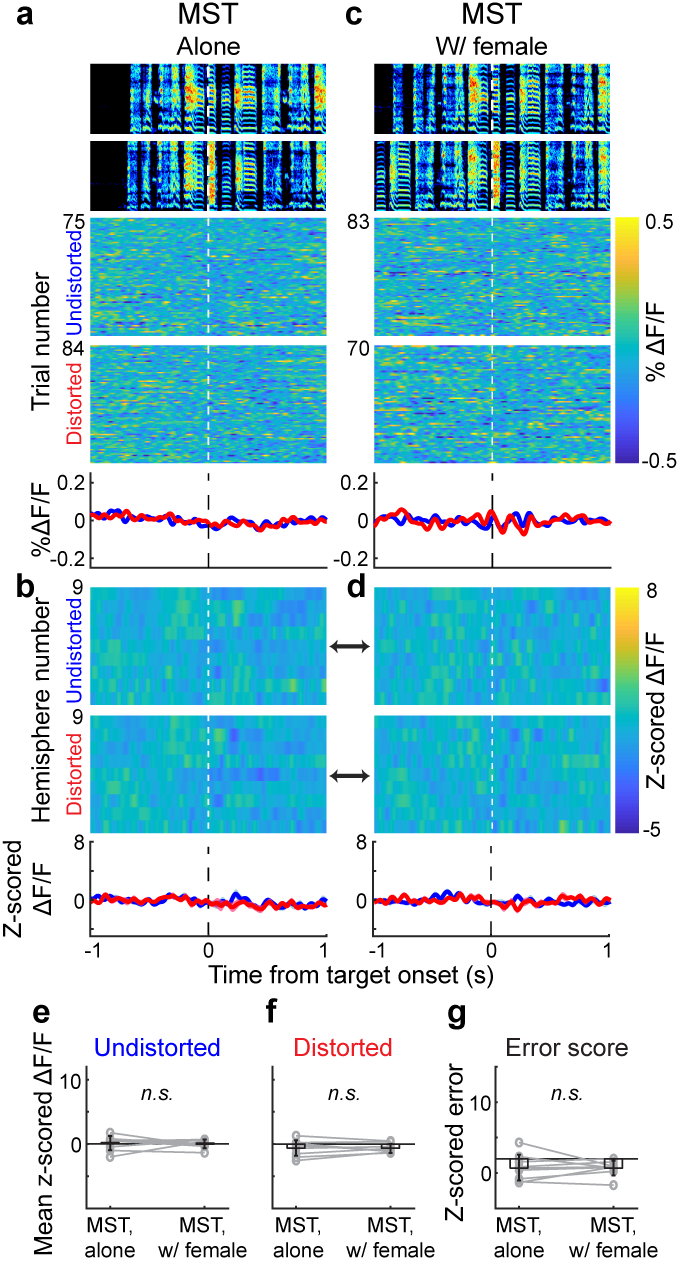
Singing-related performance error signals are not observed in MST during courtship singing. **a,** Spectrograms and single trial DA responses for undistorted (top) and distorted (bottom) renditions recorded in a single MST hemisphere of a bird singing alone, plotted above average % ΔF/F signals (different from Fig. 2c; plots aligned to target onset; blue: undistorted; red: distorted). **b,** Z-scored average from 9 MST hemispheres aligned to undistorted (top) and distorted (bottom) renditions, black arrow indicates example hemisphere shown in **a**; bottom: average z-scored response (mean ± SEM). **c,** Data plotted as in **a** for the same hemisphere measured during courtship singing. **d,** Data plotted as in **b** for the same hemispheres recorded during courtship singing. **e-f,** Scatter plots of average across all hemispheres (mean ± SD, black) and mean z-scored ΔF/F value for each hemisphere (gray) in the 150-300 ms following undistorted (**e**) and distorted (**f**) renditions in MST during alone versus female directed singing. **g,** Z-scored error responses when birds sang alone versus to females (mean ± SD, black; single hemispheres, gray). *n.s.* not significant for a 2-way ANOVA and post-hoc Tukey (**e** and **f**; Methods) and for a paired two-sided Wilcoxon signed rank test (**g**); *n* = 9 MST hemispheres.

**Extended Data Fig. 5.**
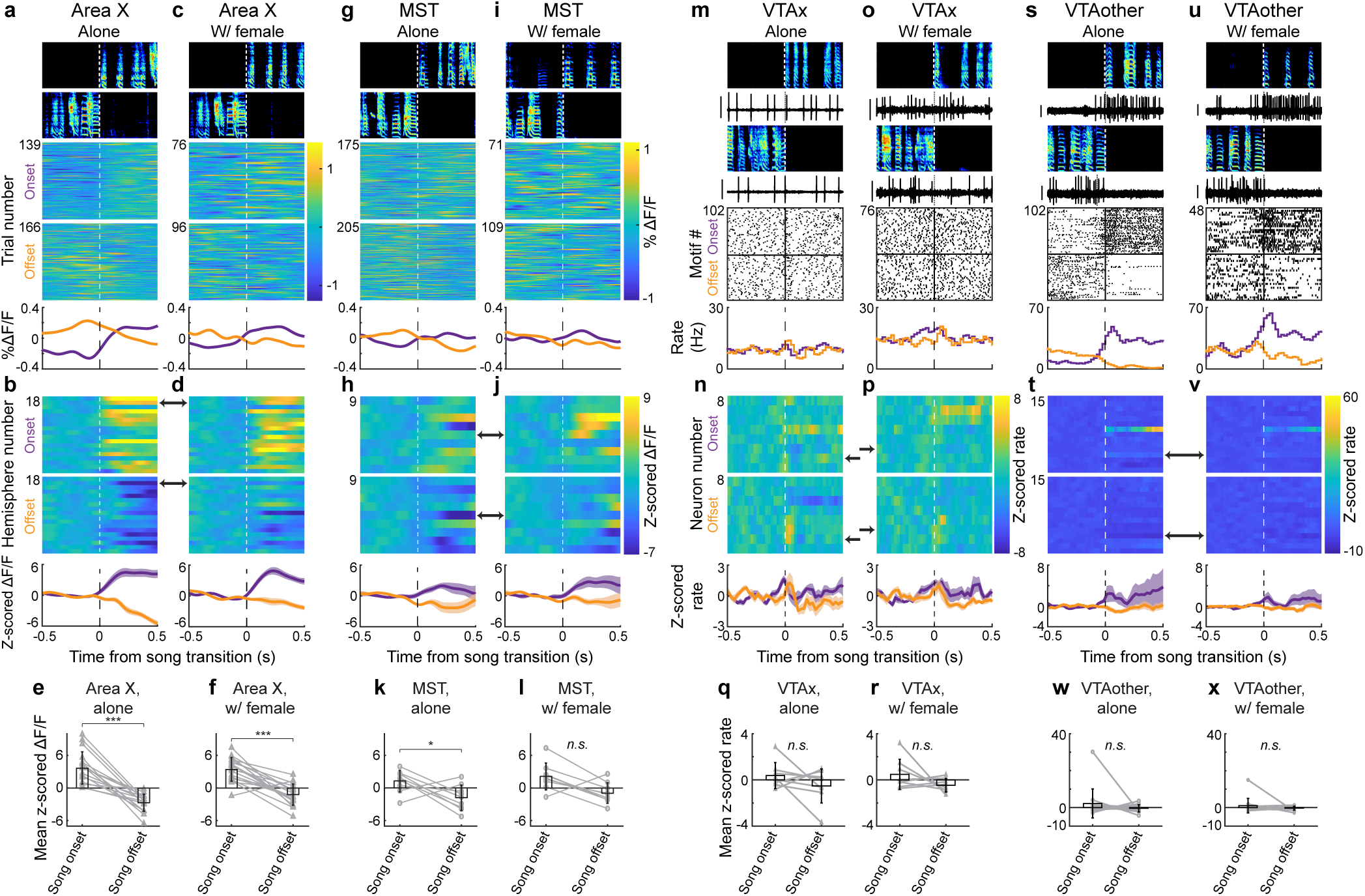
Baseline levels of DA release, but not VTA spiking, increase during both alone and female directed singing. **a,** Spectrograms and single trial DA signals for song onsets (top) and offsets (bottom) recorded in a single Area X hemisphere of a bird singing alone, plotted above average % ΔF/F signals (purple and orange aligned to song onset or offset, respectively). **b,** Z-scored average from 18 Area X hemispheres aligned to song onsets (top) and offsets (bottom), black arrow indicates example hemisphere from **a**; bottom: average z-scored signals (mean ± SEM). **c-d,** Data plotted as in **a-b** for the same hemisphere measured during courtship singing. **e-f,** Mean z-scored ΔF/F value for each hemisphere (gray) in the 0-500 ms window following song onset and offset in Area X during alone (**e**) female directed singing (**f**) (black: mean ± SD across all hemispheres). **g-l,** Data plotted as in **a-f** for MST. **m,** Spectrograms, single trial spiking activity, and raster plots for song onset (top) and offset (bottom) recorded in a single VTAx neuron of a bird singing alone, plotted above firing rate histograms (plots aligned to song onset or offset). **n,** Z-scored firing rate histograms from 8 VTAx neurons aligned to song onset (top) and offset (bottom), black arrow indicates example neuron shown in **m**; bottom: average z-scored firing rate (mean ± SEM). **o-p,** Data plotted as in **m-n** for the same VTAx neurons recorded during courtship singing. **q-r,** Scatter plots (gray) and mean ± SD (black) of average z- scored firing rate in the 0-500 ms window following song onset and offset during alone (**q**) and female directed singing (**r**) across all VTAx neurons. **s-x,** Data plotted as in **m-r** for VTAother neurons. Scale bars in **m, s** and **u** for spiking activity is 1 mV, and for **o** is 0.25 mV. * *P*<0.05, *** *P*<0.001, *n.s.* not significant for a paired two-sided Wilcoxon signed rank test; *n* = 18 Area X and *n* = 9 MST hemispheres (**e,f,k,l**) and *n* = 8 VTAx and *n* = 14 VTAother neurons (**q,r,w,x**).

**Extended Data Fig. 6.**
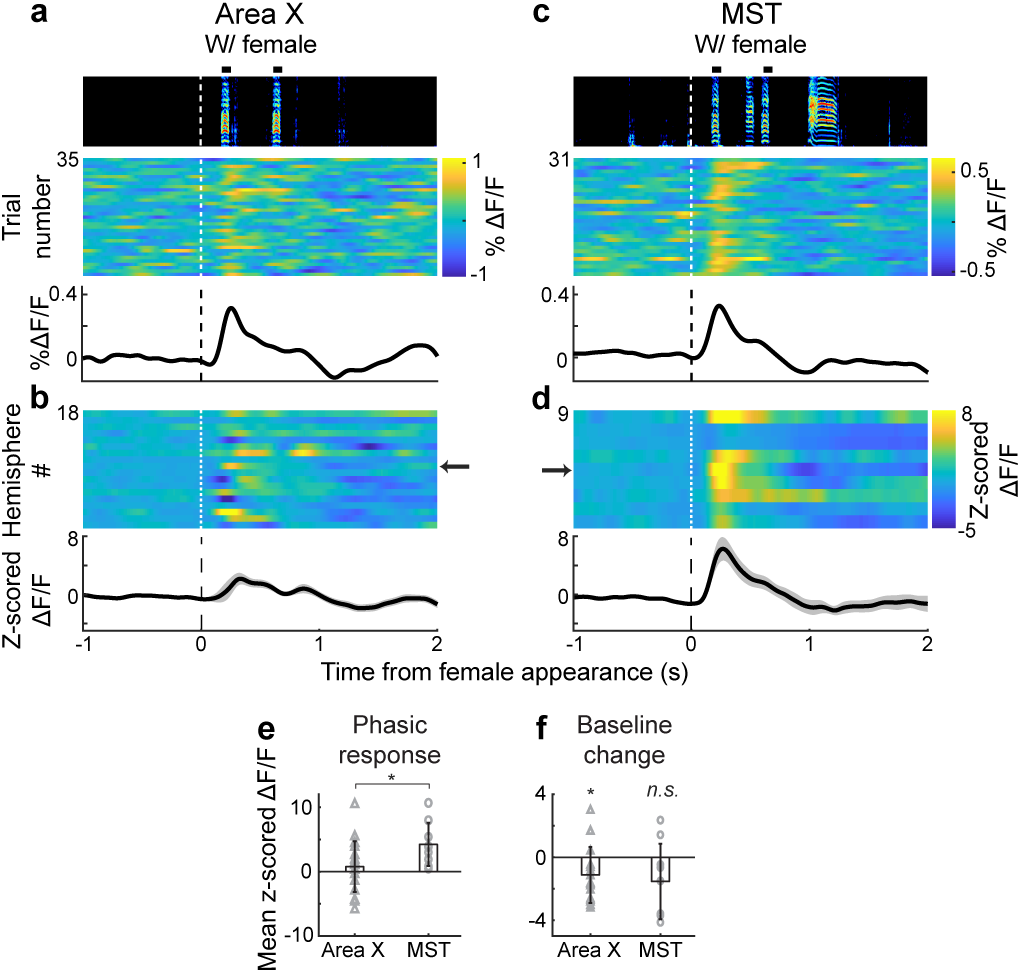
The cued moment of female appearance evokes phasic, but not sustained, activation of DA release. **a,** Spectrogram and single trial Area X DA responses to female appearance during the transition between alone and with-female conditions, plotted above average % ΔF/F signals (plots aligned to onset of female call playback). Black boxes above spectrogram indicate female call playbacks. **b,** Z-scored average from 18 Area X hemispheres, black arrow indicates example hemisphere shown in **a**; bottom: average z-scored response (mean ± SEM). **c-d,** Data plotted as in **a-b** for DA signals recorded in MST. **e,** Scatter plots showing phasic response across all hemispheres (mean ± SD, black) and mean z-scored ΔF/F value for each hemisphere (gray) in the 150-300 ms window following female call onset cueing female appearance (triangles: Area X; circles: MST). **f,** Scatter plots of baseline change across all hemispheres (mean ± SD, black) and mean z-scored ΔF/F value for each hemisphere (gray) in the 1-2 second window following cued female appearance (triangles: Area X; circles: MST) (Methods). \**P*<0.05, *n.s.* not significant for an unpaired two-sided Wilcoxon rank sum test; *n* = 18 Area X and *n* = 9 MST hemispheres (**e**), and for a one-sample t test (**f**).

**Extended Data Fig. 7.**
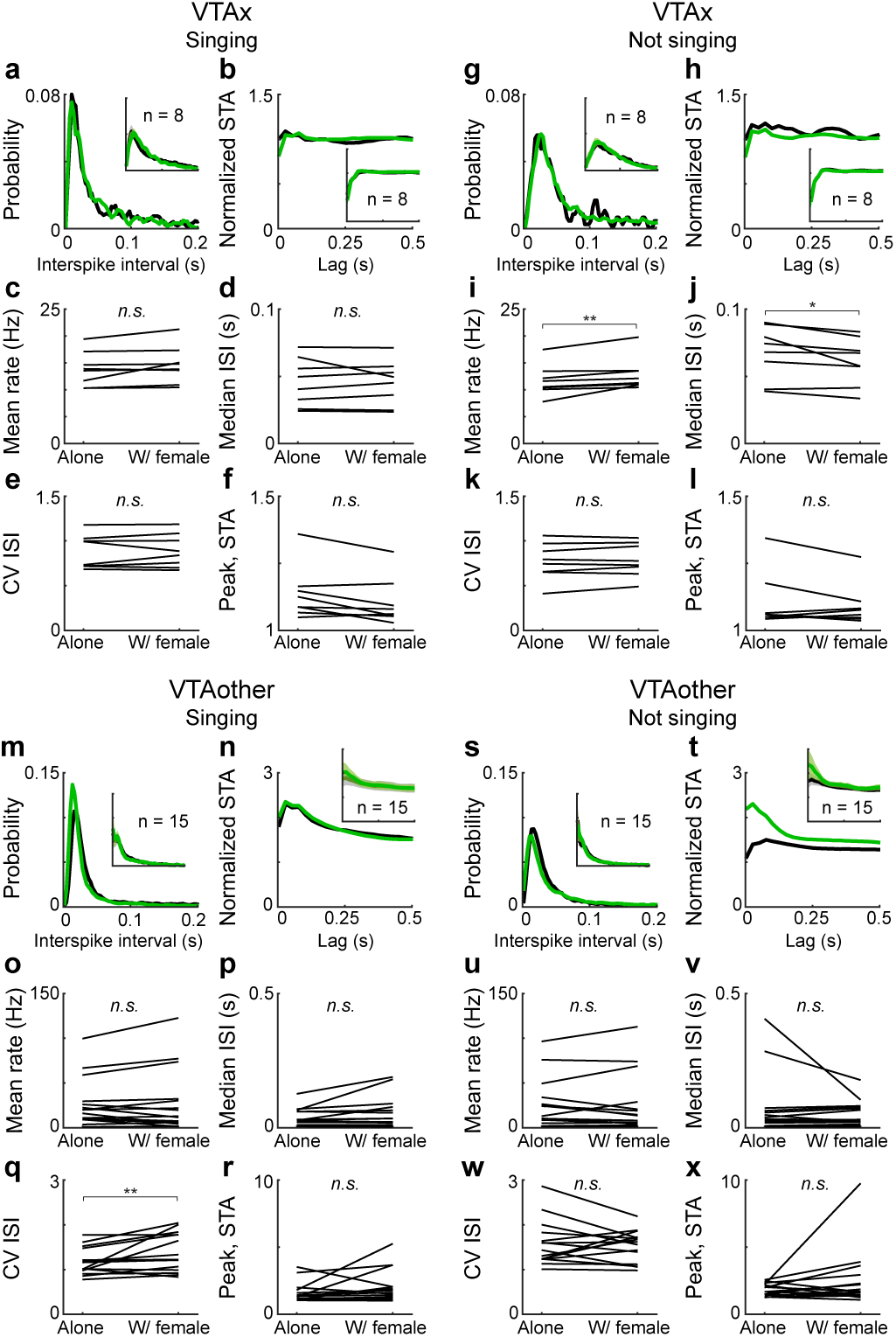
Discharge statistics of VTAx DA neurons do not depend on courtship context during singing and were subtly affected during not-singing periods. a-b, ISI distribution (**a**) and normalized spike train autocorrelogram (STA) (**b**) during singing alone (black) and female directed (green) song for a single VTAx neuron. Insets: mean ± SEM for 8 VTAx neurons. **c-f,** Mean firing rate (**c**), median ISI (**d**), coefficient of variation of the ISI distribution (CV ISI; **e**), and peak of the STA (**f**) for 8 VTAx neurons recorded when males sang alone and sang female-directed song. **g-l,** Data plotted as in **a-f** for the same 8 VTAx neurons during not singing periods. **m-x,** Data plotted as in **a-l** for 15 VTAother neurons (Methods). * *P*<0.05, ** *P*<0.01, n.s not significant for a paired two-sided Wilcoxon signed rank test.

**Extended Data Fig. 8.**
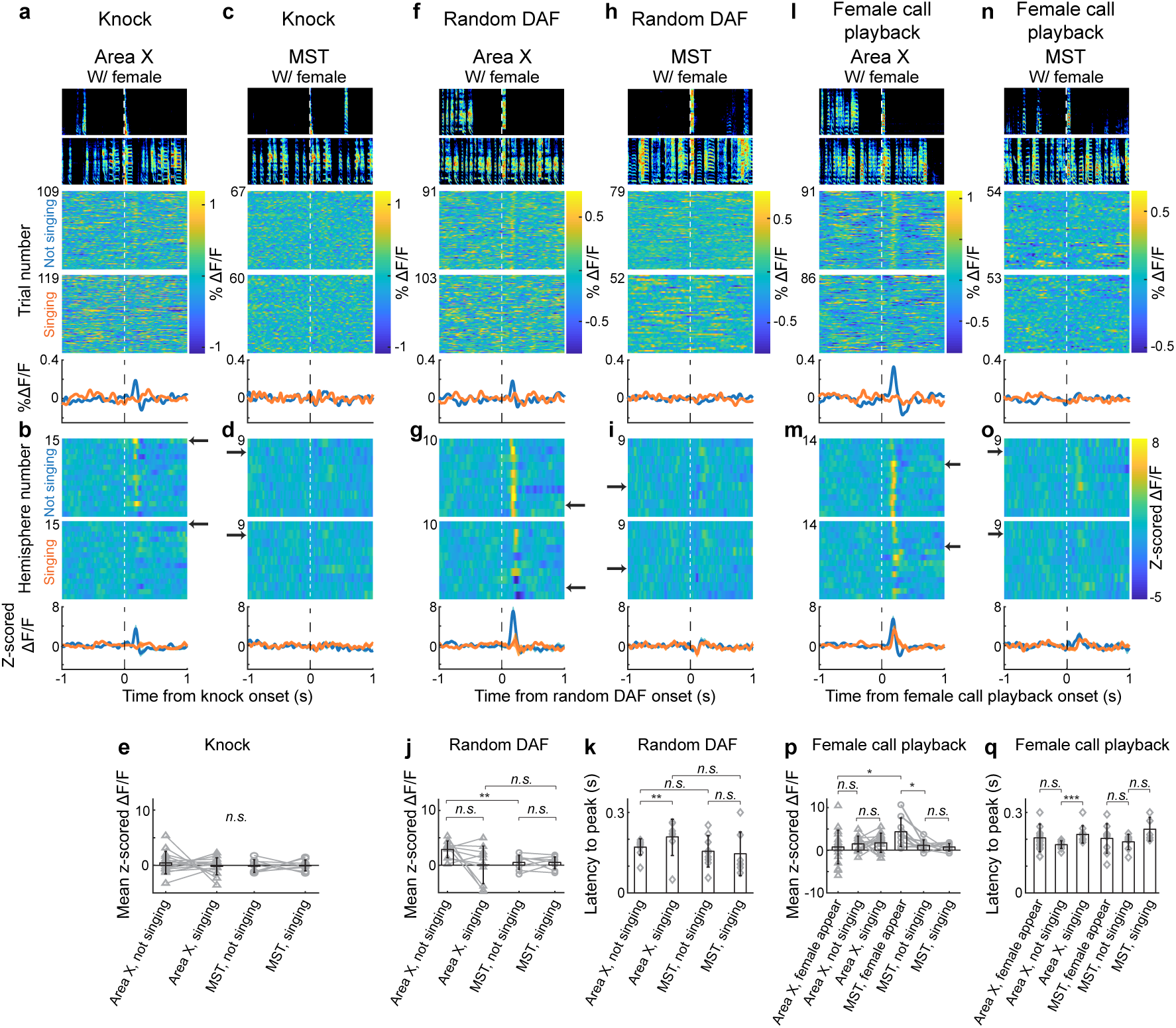
Socially irrelevant sounds and pre-recorded female calls played during courtship do not drive reliable DA responses. a-e, Knock responses. **a,** Spectrograms and single trial Area X DA responses to knock sounds played during not-singing (top) or singing (bottom) periods of the courtship interaction, plotted above average % ΔF/F signals (plots aligned to onset of knock). **b,** Z-scored average from 15 Area X hemispheres, black arrow indicates example hemisphere shown in **a**; bottom: average z-scored response (mean ± SEM). **c-d,** Data plotted as in **a-b** for knock responses recorded in MST. **e,** Mean z-scored ΔF/F values in the 0.15-0.3 second window following the onset of knock sounds in Area X and MST during singing and not-singing periods (gray, single hemispheres; black, mean ± SD across hemispheres; triangles: Area X; circles: MST). **f-k,** Random DAF responses. **f-j,** Data plotted as in **a-e** for Area X and MST responses to the sound of DAF played randomly throughout the courtship interaction, including during singing and not-singing periods. **k,** Latencies to peak responses to random DAF (gray, single hemispheres; black, mean ± SD across hemispheres; Methods). **l-q,** Pre-recorded female calls. Data plotted as in **f-k** for Area X and MST responses to the sound of pre-recorded female calls played through a speaker at random times throughout a live courtship interaction, including during singing and not-singing periods. Note that playback of these calls did not evoke reliable DA responses in Area X or MST. * *P*<0.05, ** *P*<0.01, *** *P*<0.001, *n.s.* not significant for a paired two-sided Wilcoxon signed rank test (within Area X and within MST **e,j,p**) and for an unpaired two-sided Wilcoxon rank sum test (**k, q**; across Area X and MST in **e,j,p**).

